# Individual identity and environmental conditions explain different aspects of sleep behaviour in wild boar

**DOI:** 10.1101/2022.11.23.517569

**Authors:** Euan Mortlock, Václav Silovský, Justine Güldenpfennig, Monika Faltusová, Astrid Olejarz, Luca Börger, Miloš Ježek, Dómhnall J Jennings, Isabella Capellini

## Abstract

Sleep is a fundamental behaviour as it serves vital physiological functions, yet how the sleep of wild animals is constrained by environmental conditions is poorly understood. Using non-invasive multi-sensor high-resolution biologgers and a robust classification approach, we quantified multiple dimensions of sleep in wild boar (*Sus scrofa*), a nocturnally active mammal, monitored for up to a full annual cycle. In support of the hypothesis that environmental conditions determining thermoregulatory challenges regulate sleep, we show that on warmer, longer, and more humid days sleep quality and quantity are reduced, whilst greater snow cover and rainfall promote sleep quality. Importantly, our study reveals large inter-and intra-individual variation in sleep durations, suggestive of pace-of-life syndromes. Given the major role that sleep plays in health, our results suggest that global warming and the associated increase in extreme climatic events are likely to negatively impact sleep, and consequently health in wildlife, particularly in nocturnal animals.

## Introduction

Sleep is a behaviour, observed in virtually all animals (Anafi *et al.* 2019), where individuals enter a state of quiescence in a species-specific posture and require a stronger stimulus to elicit a response compared to individuals that are in wakeful rest (Cirelli & Tononi 2008). Unlike torpor and hibernation, sleep is also characterised by a rapid return to the waking state (Tobler 2011). Sleep is beneficial, being associated with many vital physiological functions. Sleep boosts the immune system (Opp & Krueger 2015; Besedovsky *et al.* 2019), promotes endocrine production and metabolic regulation (Spiegel & Leproult 1999; Leproult & Van Cauter 2010; Morris *et al.* 2012; Manzar *et al.* 2014; Medic *et al.* 2017), and supports neural maintenance and cognitive functions, such as memory consolidation (Walker 2009; Lim & Dinges 2010; Xie 2013; Dzierzewski *et al.* 2018; Klinzing *et al.* 2019). The importance of sleep for the brain and body is further highlighted by the detrimental health- and cognition-related consequences of sleep loss (sleep deprivation) in both the short and long-term; likely to mitigate the costs of sleep loss, sleep deprivation is often followed by longer sleep (sleep rebound) (Kushida 2004). However, sleep has inherent opportunity costs since sleeping animals cannot engage in fitness enhancing behaviours like foraging or finding mates and is likely associated with greater risk of predation (Capellini *et al.* 2009). Consistently, phylogenetic comparative studies using sleep data from laboratory animals demonstrate that sleep durations and patterns are influenced by species’ ecology (e.g. Capellini et al. 2008). Sleep has so far been studied primarily in the laboratory where animals do not experience any of the benefits and costs of sleep loss. Thus, we know very little about how animals meet their sleep need, and how ecological conditions constrain sleep, in the wild.

Environmental conditions affect both the quantity and quality of sleep in the laboratory (Harding *et al.* 2020). Light and ambient temperature are well known to influence sleep in humans and animals, hence they are finely controlled in laboratory studies (Lan *et al.* 2017; Reinhardt 2020). Specifically, the first stage of deep sleep in mammals, NREM (non-rapid-eye-movement) sleep, is characterised by low, constant body temperature; a cool ambient temperature thus promotes the onset of sleep, greater sleep efficiency and quality (reduced fragmentation into multiple sleep bouts and longer sleep bouts) and total sleep duration in the laboratory (Troynikov *et al.* 2018; Harding *et al.* 2020). Conversely, the thermoregulatory challenge presented by high temperature reduces sleep time, increases sleep fragmentation and reduces sleep quality, and upregulates behaviours that help thermoregulation (Downs *et al.* 2015; Harding *et al.* 2020). Altogether, this evidence suggests that environmental conditions, such as daily weather and seasonal changes, should affect sleep in the wild. Consistently, the few studies in the wild find that temperature affects sleep in natural environments; high temperature increases time invested in licking, a thermoregulatory behaviour, at the expense of sleep in fruit bats (*Epomophorus wahlbergi,* Downs *et al.* 2015); king penguins (*Aptenodytes patagonicus)* sleep less in hotter summer days (Dewasmes *et al.* 2001); and gibbons’ sleep becomes more fragmented at higher temperatures (*Hylobates moloch/pileatus)* (Reyes *et al.* 2021). However, ambient temperature does not appear to influence sleep duration or fragmentation in baboons (*Papio anubis)* (Loftus *et al.* 2022), although this conclusion may be premature and due to the limited temperature fluctuation over the month-long recording period of this study.

Beyond ambient temperature, wild animals are exposed to many environmental conditions that change throughout the day and across the year. Humidity can compound the effects of higher temperatures on sleep, making thermoregulation more difficult by reducing the efficiency of evaporative cooling (Harding *et al.* 2019; Mota-Rojas *et al.* 2020). Thus, higher humidity should lead to shorter and more fragmented sleep. As expected, higher humidity reduces sleep duration in chimpanzees (*Pan troglodytes)* and increases sleep fragmentation in both chimpanzees (Videan 2006) and gibbons (Reyes *et al.* 2021). Conversely, rainfall and snow may promote sleep, by providing evaporative cooling or increasing the thermal value of bedding sites respectively, although we highlight that the influence of rainfall and snow on sleep in wild animals has not been studied (but see Wada *et al.* 2007)

Finally, it is well-established that light, and so day length, regulates circadian rhythm, hence when and how long to sleep (LeGates *et al.* 2014; Yadav *et al.* 2022). Hence, longer day lengths reduce, whilst shorter day lengths increase, sleep time in humans (Friborg *et al.* 2012; Yetish *et al.* 2015). Similarly, sleep is regulated by sunrise and sunset times in the nocturnal slow loris (*Nycticebus javanicus,* Reinhardt *et al.* 2019). In the wild, sleep timing and duration should thus fluctuate with changing day lengths, where longer days reduce sleep in diurnal species and increase sleep in nocturnal species. Further, if light is a cue for sleep or waking, bright moonlight may interfere with sleep regulation. Consistently, greater illumination from moonlight increases sleep duration in gibbons and humans, although moonlight does not alter sleep in baboons (Samson *et al.* 2018; Reyes *et al.* 2021; Loftus *et al.* 2022).

Importantly, while some studies in wild animals have found limited evidence that environmental conditions affect sleep time and patterns, we still do not know how sleep changes with daily and seasonal environmental variation over the annual cycle. With rare exceptions (Loftus *et al.* 2022), the few published studies on sleep in wild animals are limited by small sample sizes and short recording durations. Furthermore, some sleep studies in wild animals employed invasive recording equipment that requires surgery and recapture, and thus have likely quantified sleep in stressed individuals. If we are to understand how sleep fits within the activity budget of wild animals, how it is affected by the environment and natural constraints, and what short- and long-term costs animals pay for sleep loss, we need to study sleep in wild individuals, non-invasively, and for extended periods. Recent advances in biologging technology and analysis methods offer an ideal solution as they allow recording behaviours accurately, non-invasively, without direct observations and for long time periods in the wild (Wilson *et al.* 2008; Williams *et al.* 2020). Here, we investigate how ambient temperature, humidity, rainfall, snow, day length, and moonlight, influence sleep time, fragmentation and quality over the annual cycle in wild boar (*Sus scrofa)* that experience a broad range of environmental conditions in the wild. The wild boar is a generalist species that exhibits considerable behavioural plasticity under varying conditions (Podgórski *et al.* 2013), thus it is a good model for investigating how environmental changes influence sleep. Importantly, laboratory studies with electroencephalogram (EEG) on sleep in pigs, the domesticated relatives of wild boars (Allwin *et al.* 2016), provide valuable robust information on which to base the classification of sleep with biologgers.

Using Daily Diaries (DDs, Wildbyte Technologies Ltd), multi-sensor biologgers that allow discrimination of complex behaviour in wild animals (Wilson *et al.* 2008), we estimated total daily sleep time (TST, hours), the number of sleep bouts per day (sleep fragmentation/consolidation), and the duration of the longest daily sleep bout (sleep quality) for individual wild boar over the annual cycle. While total sleep time (TST) over 24hrs is an appropriate ecological estimate of sleep time in animals (Capellini *et al.* 2008), the number of sleep bouts over which TST occurs reflects sleep efficiency, since individuals that frequently wake up spend more time in transitional stages and less time in restorative deep sleep (Bonnet 2004, Capellini *et al.* 2009). Finally, the duration of the longest daily sleep bout in a 24-hour period indicates sleep quality as it represents the best opportunity for an individual to accrue the benefits of the most restorative stages of deep sleep (Bonnet & Arand 2003). Combined, these three daily measures of sleep provide an ecologically meaningful assessment of sleep quantity and quality. We thus predict that TST is reduced, the number of sleep bouts/day is higher, and the duration of the longest daily bout is shorter when ambient temperature and humidity are higher. Conversely, we expect that greater rainfall and snow depth increase TST, reduce the number of bouts/day, and increase the duration of the longest bout. Finally, we predict that longer day lengths increase TST, reduce the number of bouts/day, and increase the duration of the longest bout, while greater moonlight should increase the number of bouts, reduce TST and the longest bout. Moreover, unlike previous studies, we also investigate whether, and to what extent individuals differ in sleep time and patterns, and in how their sleep changes with environmental conditions.

## Methods

### Study sites

This study took place between 05/05/19 and 01/12/21 (start-to-end 941 days), in Kostelec (Central Bohemian region; 49.96N, 14.78E) and Doupov (Karlovy Vary region; 50.24N, 13.12E) in the Czech Republic (Figure S1). Kostelec is forested suburban area near Prague open to the public; Doupov is mixed forest and hills, closed to the public with military/forestry access only.

### Procedures

We employed traps to capture, immobilise, and fit 28 adult and sub-adult wild boar (24 females and 4 males) with collars bearing biologging units. We used customized Vertex Plus collars produced by Vectronic Aerospace GmBH (https://www.vectronic-aerospace.com/, Berlin, Germany), carrying Daily Diaries (DD; Wildbyte Technologies Inc, Swansea, Wales) and a standard GPS module. DD carried a tri-axial accelerometer recording at 10Hz, data was stored on-board memory cards and downloaded on collar recovery after drop-off (Wilson *et al.* 2008, Figure S2, Supplementary Methods). The duration of recording time differed among individuals from 10 to 363 days (mean 89 days), with a population total of 2424 days of data (Figure S3).

### Ethics

This work was carried out in accordance with the guidelines of the Ministry of the Environment of the Czech Republic; the trapping and handling protocol was approved by the ethics committee of the Ministry of the Environment of the Czech Republic and carried out in accordance with the decision of the ethics committee of the Ministry of the Environment of the Czech Republic number MZP/2019/630/361. A full description of trapping, immobilization, and handling procedures is available in the Supplementary Methods.

### Classification of sleep

We derived a robust procedure to identify sleep bouts with DD data by using EEG studies of sleep in domestic pigs to precisely describe sleep postures and derive rules to identify these in the accelerometer data. These studies identify two sleep postures in pigs; lateral or sternal recumbency with the head on the ground (Ruckebusch 1972; Skinner *et al.* 1975; Kuipers & Whatson 1979), accompanied by rapid loss of muscle tone at sleep onset (Ruckebusch 1972). To derive posture from the raw DD acceleration data (in *g*) we calculated the “static acceleration”, the degree of acceleration due to gravity only (Wilson *et al.* 2008). Then, from a running mean of the static acceleration (“smoothed”, calculated over two seconds, or 20Hz) we computed the body pitch and roll angles using the arcsine of the *g* for the surge (pitch) and sway (roll) axes (e.g. 0.98*g* on the surge axis equals 0° pitch; Shepard *et al.* (2008). Pitch and roll angles were smoothed over two seconds and sternal recumbency with head-down was defined as (*pitch* < 0°) and (*roll* > −15° and +15° <), while lateral recumbency was defined as (*roll* < −15° and > +15°). Sustained lack of movement, the other key behavioural cue for sleep, was identified using VeDBA smoothed over two second. VeDBA is the sum of the vector of the dynamic acceleration (raw acceleration with the gravitational component removed), calculated as;

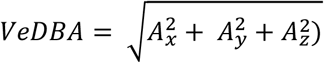

where *A_x_*, *A_y_*, *A_z_* are the dynamic components of each axis of acceleration (see Williams *et al.* 2020). We set a threshold for movement in sleep postures to 0.2 VeDBA where sleep bouts ended if this threshold was crossed, allowing small postural changes during sleep. Finally, given that domestic pigs in sleep posture require 4-5 minutes to transition from wakefulness to sleep (Ruckebusch 1972: 5 minutes 50 seconds; Robert and Dallaire 1986): 4.11 ± 3.32 minutes), we discarded the first 5 minutes as ‘transitional state’ from all periods of data where the criteria for sleep posture and lack of movement were met. This classification therefore separates sleep from wakeful rest, using the behavioural markers for sleep (Figure 1, Figures S4 & S5).

**Figure 1:**
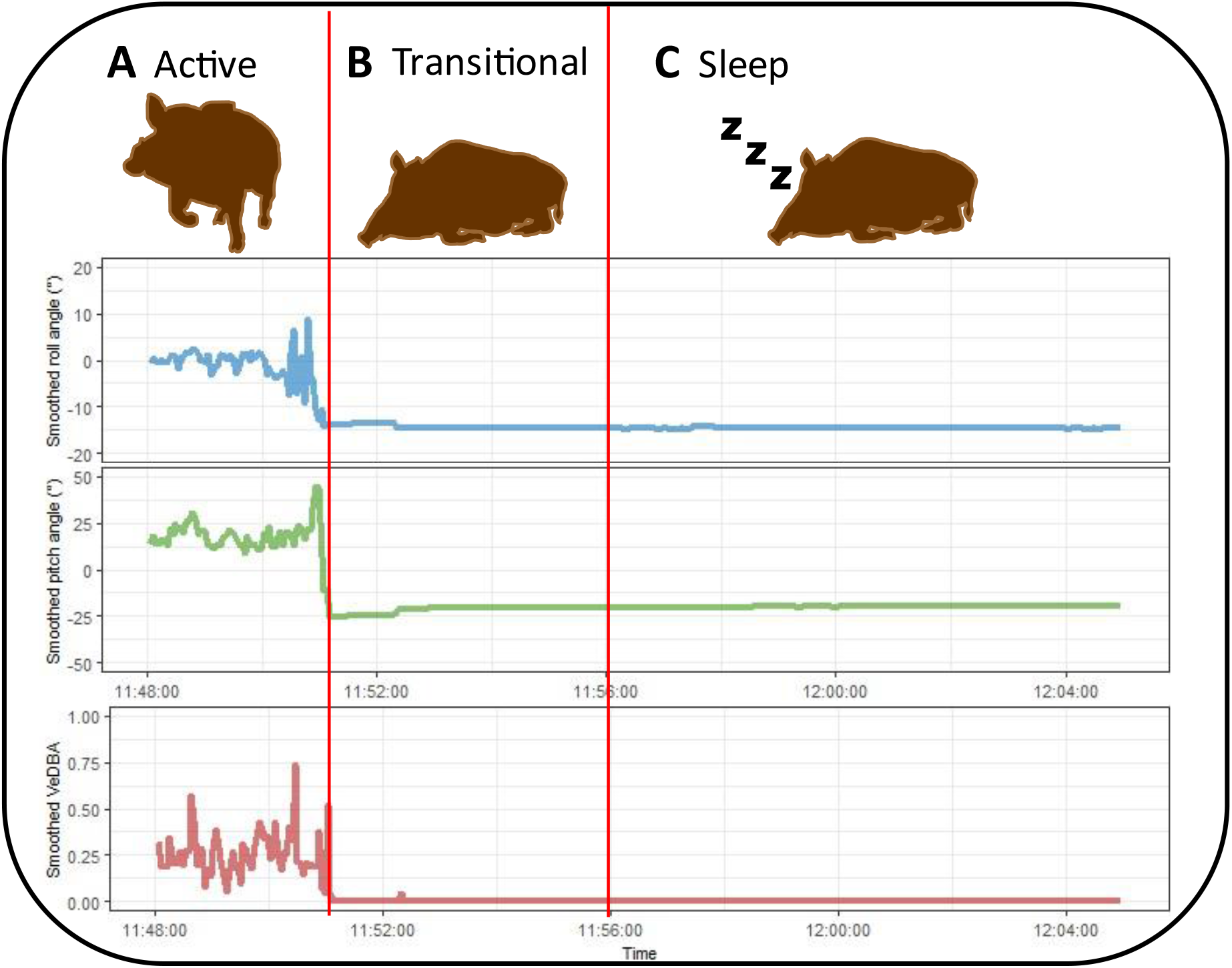
Visualisation of DD data showing changes in smoothed roll, pitch, and VeDBA values, corresponding to relevant behavioural types (separated by red lines), to identify the onset of sleep. (A), patterns typical of general active behaviours such as movement and foraging; (B) patterns typical of sternal recumbency for the period of drowsiness/transitional sleep; and (C), an individual is classified as asleep in sternal recumbency for longer than the 5-minute window with little to no movement.

When considering individual differences collected from movement data, it is necessary to carefully check that these differences do not arise from measurement error, equipment malfunction, or data processing (Hertel *et al.* 2020). In order to address this, all boar were fitted with the same devices and the data were processed in the same manner. We visually inspected processed and raw accelerometry data to ensure there were no sources of error.

### Environmental data

We drew hourly weather data from the Jevany (Kostelec, 49.96N, 14.80E) and Kyelska Spa (Doupov, 50.26N, 13.02E) weather stations (www.visualcrossing.com). Daily means were computed for ambient temperature (degrees Celsius, °C); snow depth (cm); and relative humidity (the amount of water vapor present in the air compared to the maximum amount possible for a given temperature, as a percent, %). Precipitation (mm) was quantified as the total daily precipitation. Day length (hours) was estimated as hours of light from sunrise to sunset. Moon phase was coded as a continuous variable ranging from new moon (dark; 0) to full moon (bright, 1). Table S1 reports the range of environmental conditions recorded over the study period.

### Statistical analysis

Following Hertel *et al.* (2020), we used double-hierarchical generalised linear mixed-effects models (DHGLM) to assess how wild boar altered their sleep in relation to changing environmental conditions. Specifically, we modelled the changes in the mean of the three sleep measures (TST, number of daily sleep bouts, longest sleep bout) and their variance (“sigma” component) in a Bayesian framework with the R package ‘brms’ (Bürkner 2017, 2018; R Core Team 2022; RStudio Team 2022), and the Stan open source modelling platform (Stan Development 2022). Unlike standard linear mixed-effects models, DHGLM models can handle non-heterogenous residual errors, allowing a more robust assessment of fixed effects (Bridger *et al.* 2015), which is thus suitable for data of different individuals recorded over time (Figure S2).

Prior to analyses, the longest bout/day was log-transformed and all fixed effects were centred and scaled (Kruschke 2015). We assigned Gaussian distributions to response variables. We included an autoregression term of order 1, applied to each individual, to control for temporal autocorrelation. As our data structure was hierarchical, where measures of sleep were nested inside individual ID, we included ID as a random effect for both the mean and sigma component of each model, to determine whether inter- and intra-individual variation in the three sleep measures varied by individual. We controlled for location, sex, and year of data collection by including these as fixed effects in model.

We ran models using Markov chain Monte Carlo (MCMC) with weakly informative, normally distributed priors for the fixed effects (for TST and number of bouts/day models: mean of zero and a variance of 10; for longest bout/day model: mean of zero and variance of 100). We assigned weakly informative, scaled t-distributed priors with 3 degrees of freedom (Gelman *et al.* 2008) to the random effects (individual-level variation) and error terms in both components of the models. We ran chains of 15,000 iterations with a burn-in of 1,000 iterations for TST and number of bouts models, and 30,000 iterations with a burn-in of 20,000 iterations for the longest bout model, sampling every 15^th^ iteration. Visual inspection of the traces in the resulting posterior distributions showed adequate mixing and convergence. The Gelman-Rubin convergence statistic (Rhat) showed satisfactory convergence as values were equal to 1 for all parameters (Gelman *et al.* 2013). Effective sample size (ESS) for all estimated parameters over 1000 confirmed that the posterior distributions had negligible levels of autocorrelation (Tables S2-4). Models were run in triplicate and converged on qualitatively similar solutions.

Environmental, sex, and location variables were entered simultaneously as predictors in a starting ‘maximal model’ and treated as fixed effects. We used a model reduction approach to identify meaningful predictors (Crawley 2012). Thus, from ‘maximal models’ with all fixed predictors we removed the least meaningful predictor, re-ran the model and repeated the procedure until only meaningful predictors remained in a minimal statistically justifiable model (‘reduced models’). Predictors were classed as meaningful if the percentage of their posterior distribution crossing zero in the opposite direction of the effect was less than 5 (percentage cross-zero: P_x_; e.g. Capellini *et al.* (2015). Models also included month of the year to account for seasonal changes not captured by environmental predictors. Because the effect of “month” is cyclical (e.g. where month 12 is more similar to month 1 than month 6) we used a nonlinear second-order polynomial term. This was applied both as a fixed effect and a random slope term in the model formula. We used the Widely Applicable Information Criterion (WAIC) to confirm that models fitted with the random slope for month provided a better fit to the data than an intercept-only model (□WAIC > 7 indicates a superior model fit).

From the model random effects, we used the individual-level mean variance of each sleep measure, and its residual variance to calculate residual intra-individual variation (rIIV), i.e. how predictable each individual was in sleep. We then calculated the coefficient of variation in predictability (CVp); a measure of among-individual variation in predictability, standardised and comparable across studies (Cleasby *et al.* 2015). CVp closer to 0 indicates a population of more predictable individuals and a CVp closer to 1 indicates a population of less predictable individuals.

## Results

### Descriptive statistics

Across the study period boars slept on average for 10.6 hours/day (mean TST, SD ±3.4 hours) divided in 21 sleep bouts (mean, SD ±40 bouts) averaging 31.4 minutes (mean, SD ±40.8 minutes); mean longest sleep bout was 2.5 hours (SD ±1.38). All sleep parameters showed qualitative inter-individual variation, e.g. the shortest-sleeping individual slept for 6.4 hours per day on average; longest-sleeper slept for 14.8 hours. Most sleep occurred during the early morning and middle of the day (Figure S6), with the longest sleep bout usually beginning at 0400 or around 1200.

### Total Sleep Time (TST, hours)

From a maximal model with all predictors, the reduced model showed that TST was shorter with higher temperature (median [95% CI]: −0.55 [−0.76, −0.36]) and humidity (−0.12 [−0.24, −0.01]), longer day length (−0.22 [−0.41, −0.04]), and fuller moon phase (−0.14 [−0.24, −0.04], Figure 2, Table S2). Furthermore, individuals at Kostolec slept more daily than those at Doupov (1.96 [0.27, 3.43]), and boars slept less in 2020 and 2021 than 2019 (2020: −3.34 [−4.88, −1.90], 2021: −4.70 [−5.94, −3.42]; Figure 2, Table S2). The random effects of the model showed that individual boar differed in their mean TST (1.23 [0.85, 1.86]) where model-derived estimates varied from 12.90 to 16.66 hours across individuals (Table S2, Figure 3A). Individual boar also differed in their variance (rIIV, 0.24 [0.17, 0.33]), where model-derived estimates varied from 1.23 to 2.72 hours across individuals (Table S2, Figure 3B) with a CVp of 0.24 [0.17, 0.33]. The individual-level model estimates showed no correlation between mean TST and variance in TST (0.11 [−0.32, 0.52], Table S2).

**Figure 2:**
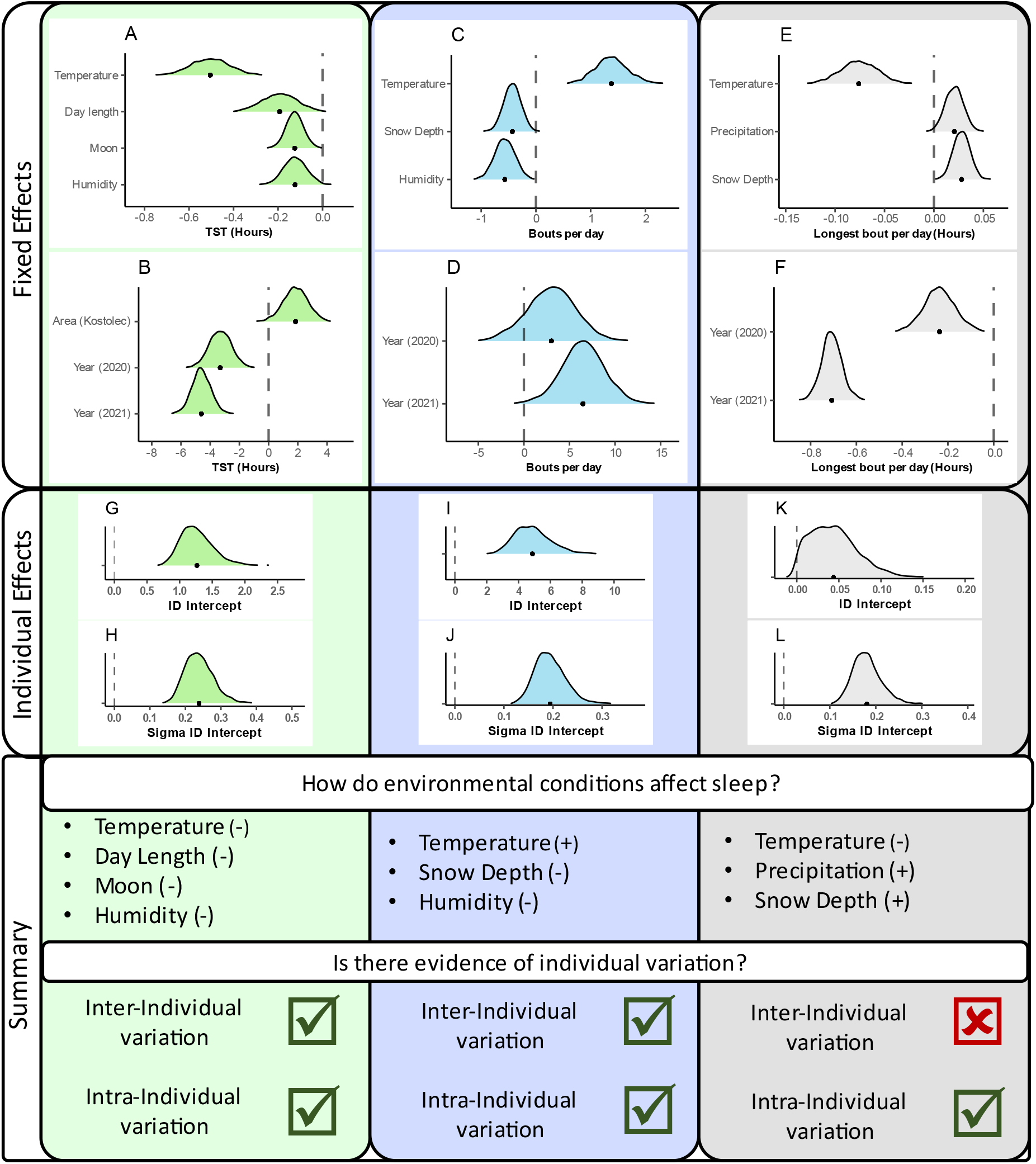
Reduced model results for total sleep time (TST, green), number of bouts/day (blue), and duration of the longest sleep bout/day (grey). (A-F); the posterior distributions for the fixed effects of environment, year, and location variables, with dashed line denoting 0. (G, I, K); posterior distributions of the random effect intercept for the mean (inter-individual variance), and (H, J, L) the posterior distribution of the random effect intercept for the sigma (intra-individual variance) component for individual ID (individual effects). Direction of effect for environmental variables (denoted with +/-), and evidence of individual variation from the individual effects, are summarised in the Summary panel (Summary).

**Figure 3:**
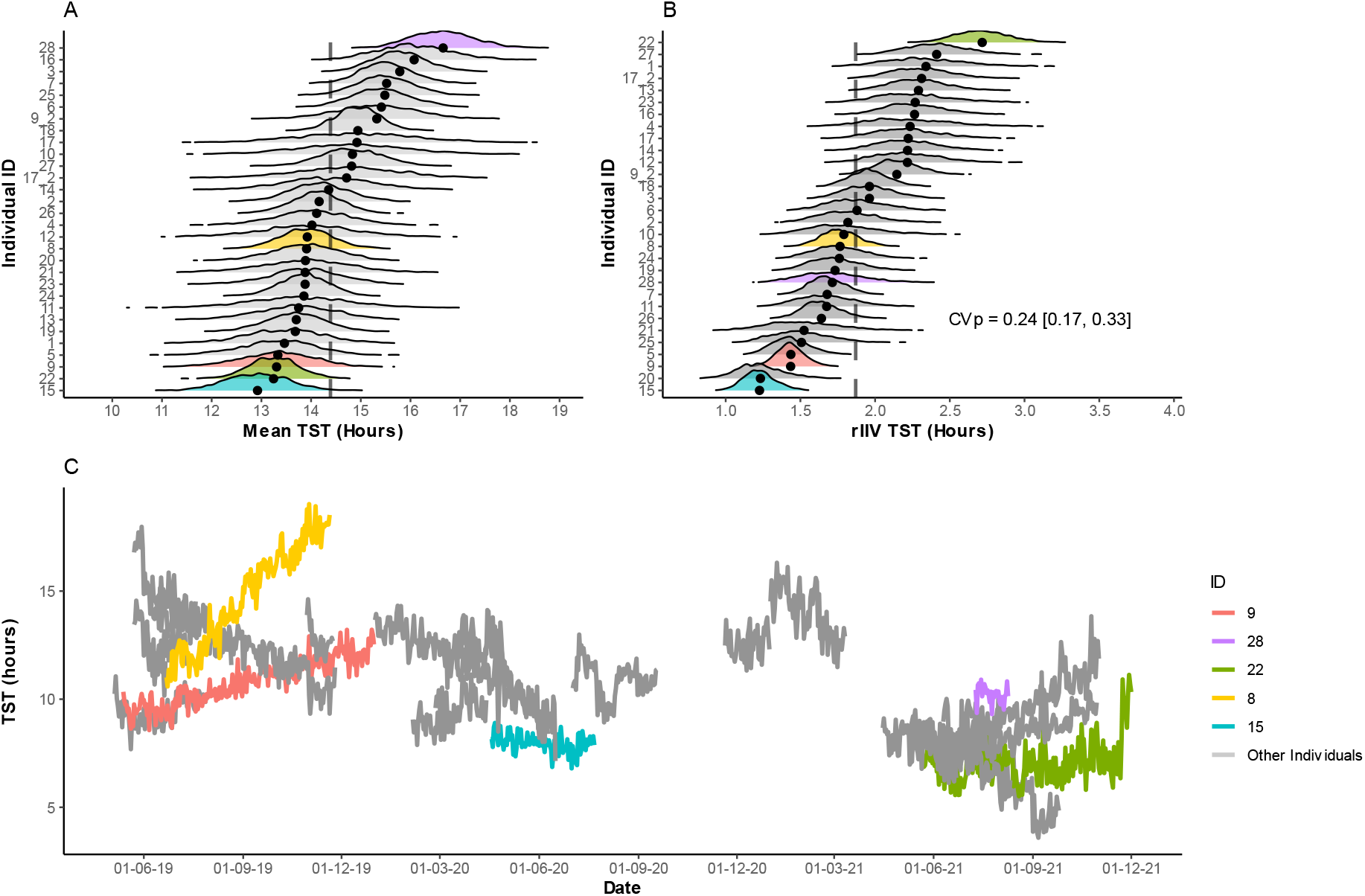
Random effects for individual ID from the reduced model of TST. A; posterior densities of mean TST estimates by individual, where points denote the individual mean, and dashed line denotes population median. Selected individuals coloured to illustrate more extreme and average individuals; the colouration is maintained through figures 3-5. B; posterior densities of TST residual intra-individual variation (rIIV), where points denote the individual mean, dashed line denotes population median, and grey shading denotes population 95% credible interval. C; model estimated values for TST for all individuals across the recording period to visualise temporal variation at the individual level and discrimination of inter- and intra-individual variation.

### Number of sleep bouts/day

From a maximal model with all predictors, the reduce model found that the number of sleep bouts/day increased with temperature (1.31 [0.59, 2.05]), and declined with greater humidity (−0.78 [−1.14, −0.40]) and snow depth (−0.44 [−0.84, −0.03]) (Figure 2, Table S3). In addition, boar slept in more bouts/day in 2021 than 2020 (2020: 2.72 [−2.58, 7.58]; 2021: 6.14 [1.72, 10.44]; Figure 2, Table S3) or 2019 (reference level). The random effects of the model revealed that individual boar differed in the mean number of sleep bouts/day (4.78 [2.97, 7.35]) where model estimates varied from 11.46 to 28.85 bouts across individuals (Table S3, Figure 4A). Individual boar also differed in their variance (rIIV, 0.19 [0.14, 0.26], Table S3, Figure 4B), where model estimates varied from 3.89 to 9.60 bouts/day across individuals (Figure 4B), with a CVp of 0.20 [0.14, 0.26]. The individual-level model estimates showed a positive correlation between mean number of sleep bouts/day and variance in the number of sleep bouts/day (0.54 [0.06, 0.84], Table S3), indicating that boar that slept in more bouts exhibited a higher variance in the number of bouts.

**Figure 4:**
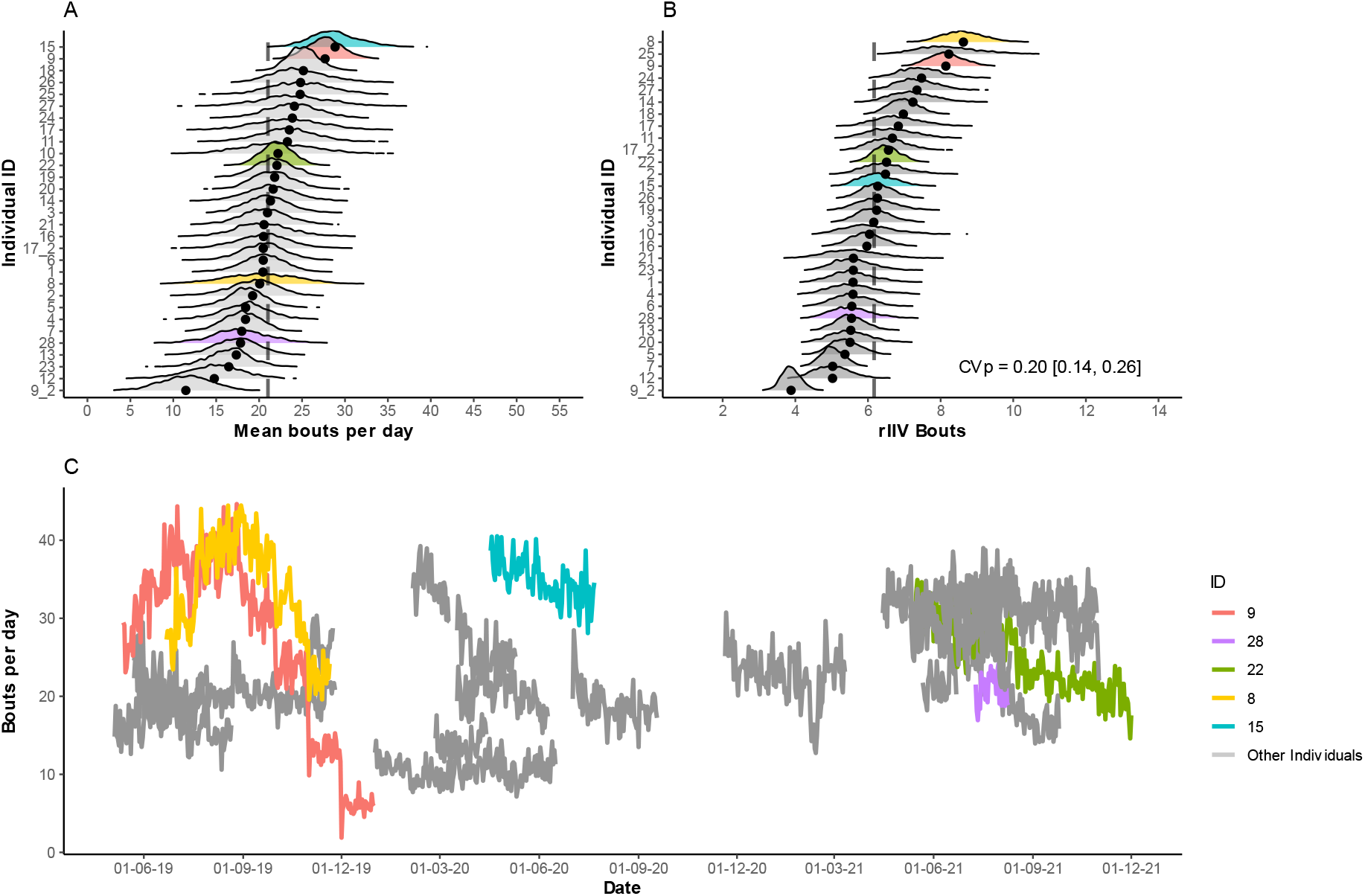
Random effects for individual ID from the reduced model of bouts/day. A; posterior densities of mean bouts/day estimates by individual, where points denote the individual mean, and dashed line denotes population median. B; posterior densities of bouts/day residual intra-individual variation (rIIV), where points denote the individual mean, dashed line denotes population median, and grey shading denotes population 95% credible interval. C; model estimated values for bouts/day for all individuals across the recording period to visualise temporal variation at the individual level and discrimination of inter- and intra-individual variation.

### Longest sleep bout

From a maximal model with all predictors, the reduced model showed that the duration of the longest bout/day decreased with increasing temperature (−0.08 [−0.11, −0.04]), and increased with greater precipitation (0.02 [0.00, 0.04]) and snow depth (0.03 [0.01, 0.04], Figure 2, Table S4). The longest sleep bout was shorter in both 2020 and 2021 compared to 2019 (2020: −0.24 [−0.37, −0.10]; 2021: −0.71 [−0.80, −0.61]; Figure 2, Table S4). The random effects of the model revealed that individual boar did not differ in the estimated mean duration of the longest sleep bout (0.04 [0.00, 0.11]) (Table S4, Figure 5A). Individual boar however differed in their variance (rIIV, 0.18 [0.13, 0.25]), where model estimates varied from 1.39 to 1.75 hours per day across individuals (Table S4, Figure 5B), with a CVp of 0.18 [0.12, 0.25]. The model estimates at the individual level showed no correlation between mean duration of the longest bout and variance in the longest bout (−0.44 [− 0.91, 0.53], Table S4), indicating that boar with greater duration for the longest sleep bout did not exhibit higher variance in its duration.

**Figure 5:**
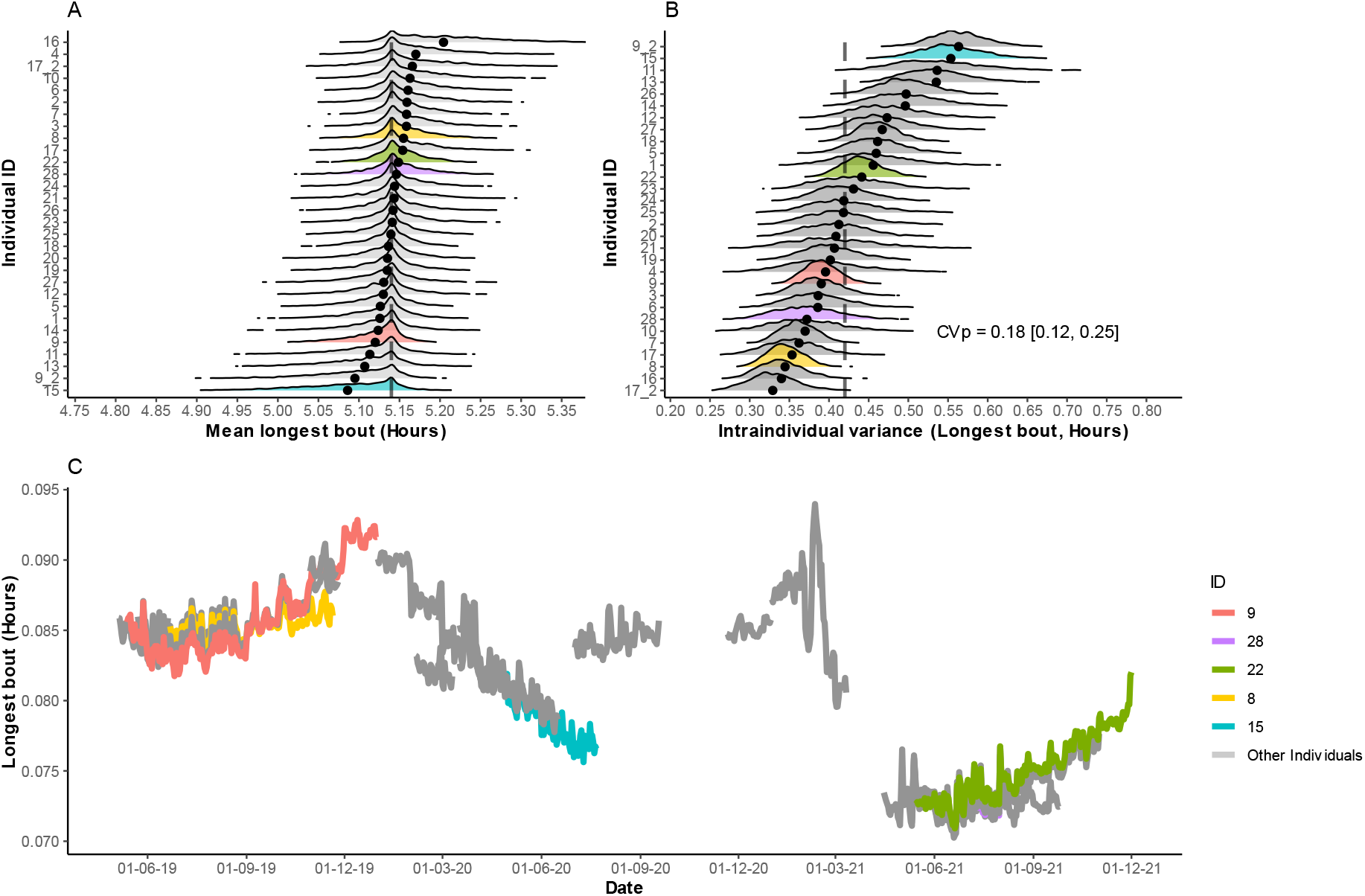
Random effects for individual ID from the reduced model of longest bout/day. A; posterior densities of mean longest bout/day estimates by individual, where points denote the individual mean, and dashed line denotes population median. B; posterior densities of longest bout/day residual intra-individual variation (rIIV), where points denote the individual mean, dashed line denotes population median, and grey shading denotes population 95% credible interval. C; model estimated values for longest bout/day for all individuals across the recording period to visualise temporal variation at the individual level and discrimination of inter- and intra-individual variation.

## Discussion

Sleep is vital, yet its patterns and tradeoffs under changing ecological conditions are largely unknown for wild animals. Investigating sleep outside the laboratory for extended periods of time is thus essential, if we are to understand its ecology and evolution. EEG studies of sleep indicate that environmental conditions related to thermoregulation and light affect sleep quantity, fragmentation and quality in the laboratory; however, their role in wild animals is poorly understood (LeGates *et al.* 2014; Harding *et al.* 2019). Using cutting-edge biologging technology we measured sleep quantity (daily total sleep time, TST), fragmentation/consolidation (number of sleep bouts/day) and quality (duration of the longest bout), in wild boar in their natural environment. Our study demonstrates that sleep quantity, fragmentation and quality varied with changes in environmental conditions and reveals that individuals differ substantially in their total daily sleep and fragmentation.

Laboratory studies have shown that environmental conditions are an important mediator of sleep behaviour (Kräuchi & Deboer 2011). The few, short-term field studies on sleep mostly confirm that temperature and light influence sleep quantity because of their effect on thermoregulation and as cue for circadian rhythms (e.g Davimes *et al.* 2018). Our study reveals that, over the annual cycle, TST in wild boar is reduced not only in warmer days but also in longer days and in more humid conditions. Hot and humid days, combined with wild boar’s preference to sleep during day time, present a major challenge for thermoregulation in this species since wild boar lack sweat glands and need to optimise body temperature by seeking out wallows (Singer *et al.* 1981). Because wallows are often located in irrigation ditches and similar areas near to human habitation, the perceived predation risk or disturbance by people is likely high and may have further detrimental impacts on sleep quantity in this species (Stuber *et al.* 2014). We further found that a more advanced moon phase reduces sleep to the same magnitude as humidity, indicating that, unexpectedly, moonlight does not favour but rather reduce sleep time in this nocturnally active species. Human disturbance is instead the probable cause of the shorter TST in 2020 and 2021, years during which human use of forests increased as a result of COVID-pandemic (A. Olejarz, personal communication Nov. 2022). Altogether, our analysis suggests that the influence of environmental conditions on TST may be exacerbated or mitigated by species traits (activity time, thermoregulatory ability) and their interaction with external natural or anthropogenic disturbance.

Although an often-overlooked aspect of sleep behaviour, sleep fragmentation (sleep that is distributed over an increasing number of bouts) is, like sleep deprivation, associated with negative effects on physiology and cognition (Stepanski 2002; Bonnet & Arand 2003; Mezick *et al.* 2009). Environmental conditions that reduce TST also increase sleep fragmentation in the laboratory (Harding *et al.* 2020). Consistently, we find that higher temperature is associated with more sleep bouts/day, hence greater fragmentation, in wild boar. Thus, a reduction in TST in warmer conditions with a concurrent increase in sleep bouts likely leads to more severe effects on health. Unexpectedly, however, sleep is less fragmented with greater humidity although this effect is small. In support of the hypothesis that snow and rainfall promote sleep quality and quantity by favouring thermoregulation through evaporative cooling or increasing the thermal value of bedding sites (Harding *et al.* 2019), sleep in wild boar is less fragmented with higher snow depth and more precipitation. Overall, we conclude that, like TST, sleep fragmentation in wild boar respond plastically to changing environmental conditions over the annual cycle. Finally, given that the number of sleep bouts was higher in 2020, we suggest that human disturbance may not only reduce time for sleep but also increase its fragmentation and ultimately have serious detrimental effects on health in wild animals.

Lastly, we investigated how the duration of the longest sleep bout responded to changing environmental conditions. Although not generally considered in sleep studies, the duration of the longest bout/day is a good indicator of sleep quality because it offers the best opportunity to gain the key benefits of the deepest and most restorative sleep stages (Bonnet 2004). As predicted, warmer temperature reduces the duration of the longest bout while greater precipitation and snow depth increase it. Similar to TST, the longest sleep bout was shorter in 2020 and 2021. Overall, sleep quality (longest bout) and quantity thus respond similarly to environmental conditions. Bringing results of the three sleep parameters together, we conclude that environmental conditions that are known to influence thermoregulation in the laboratory affect sleep in wild animals. Specifically, sleep is shorter, more fragmented and of lower quality in warmer temperature; precipitation and snow favour sleep consolidation into fewer bouts and sleep quality, while greater humidity reduces TST but this negative effect is compensated by a greater sleep consolidation. Given the complex way in which sleep behaviour responds plastically to changing environmental conditions, future studies on sleep in wild animals should thus consider more than TST for a more comprehensive understanding of how the benefits of sleep are achieved (or compromised) under natural conditions.

Our analytic approach allows us to decompose inter- and intra-individual variation (individual mean and predictability, Hertel *et al.* 2020) in sleep and reveal that individuals are distinct in daily TST and fragmentation, but are similar in the duration of the longest sleep bout. Individuals also differ in the plasticity of TST and fragmentation but not of the longest bout. Thus, all individuals appear able to satisfy a physiological minimum requirement for sleep, as quantified by the longest sleep bout. We propose that the longest sleep bout could be a better species-specific indicator of sleep need compared to the commonly estimated TST, as we find that the latter varies in both mean and variance across individuals. In support of this suggestion, we note that the variable EEG estimates of TST in domestic pigs fall somewhat in the middle of the distribution of sleep times for wild boar in this study (Ruckebusch 1972: 7.82 hours; Robert and Dallaire 1986 9.15 hours), even when considering days with similar environmental conditions to those in the laboratory (under such conditions wild boar sleep ranges from 5.0 hours to 14.7 hours, mean 10.6 hours). Consequently, differences in laboratory estimates of TST, typically based on few individuals and short recording periods, likely reflect sampling effects. Altogether, our results call into question the conclusions of some earlier studies that animals in the laboratory sleep more (or less) than in the wild.

Given the established benefits of sleep for the body and brain (Xie 2013; Opp & Krueger 2015), we propose that sleep can be viewed as a behaviour favouring self-maintenance and survival and its variation among individuals that we documented may be explained by the extended pace-of-life syndrome theory (extended POLS; Dammhahn *et al.* 2018. According to POLS, individual differences in behavioural and physiological traits covary with life history traits. Specifically, “fast-living” individuals are expected to grow quickly, invest more in reproduction and less in self-maintenance, and ultimately die younger. At the opposite extreme, “slow-living” individuals should grow slowly, invest more resources in self-maintenance than reproduction, and live longer. Thus, shorter sleep may represent a facet of the “fast” living strategy where sleep, as a self-maintenance process, is reduced in favour of behaviours that enhance reproductive investment. Consistent with this suggestion, individual-level differences in TST in fruit flies (*Drosophila melanogaster)* are genetically determined and shorter-sleeping flies die younger (Cirelli *et al.* 2005; Anderson *et al.* 2022). Given that sleep loss comes at substantial costs (Rechtschaffen *et al.* 1989; Bonnet & Arand 2003; Kushida 2004; Guyon *et al.* 2014), we thus expect that individuals with shorter and more fragmented TST exhibit reduced immunocompetency and impaired cognitive abilities such as decision making (e.g. response time to approaching predators). Future research should investigate whether short-sleeping individuals within-species show correlated tendencies with traits such as growth rate and reproductive behaviour.

To conclude, our study demonstrates that, over the annual cycle, sleep in the wild is shaped by changing environmental conditions that affect thermoregulation. Importantly, sleep is shorter, more fragmented and of lower quality in warmer temperatures. Our study also reveals profound individual level differences in daily sleep quantity and efficiency, and in plasticity. Given the major role that sleep plays in health (Klingenberg *et al.* 2012; Chaput *et al.* 2017; Besedovsky *et al.* 2019), global warming and the associated increase in extreme climatic events, are likely to negatively impact sleep, and consequently health, in wildlife, particularly in nocturnal animals. Such detrimental effects may be further exacerbated if wild animals are exposed to anthropogenic stressors that disrupts sleep.

## Acknowledgements

EM is funded by the Department for the Economy (DfE), Northern Ireland. This work was supported by EVA4.0, no. CZ.02.1.01/0.0/0.0/16_019/0000803 financed by OP RDE, supported by grant of Ministry of agriculture of Czech Republic no. QK1910462 to MJ.

