## Supplementary Information for "Individual identity and environmental conditions explain different aspects of sleep behaviour in wild boar"

### Supplementary Methods

#### Trapping protocol

Boar were captured using wooden corral traps which are baited on a regular basis with corn. The trap is triggered when a boar enters the trap and disturbs a weight, releasing the door trigger mechanism and allowing the corral door to slide down and close. After the door trigger mechanism is released, an alarm device remotely notifies the handlers responsible for the collaring. Handlers stationed nearby then anesthetized trapped boar using a mixture of Tiletamin, Zolazepam, Ketamin and Xylazin in narcotization darts delivered via airgun (for details see Barasona *et al.* (2013)). During the handling process, which took approximately 20 minutes, physical measurements and tissue samples were taken as part of a broader study. The trapped individual was marked with a plastic ear tag and lastly fitted with a collar. Once handling was completed, handlers withdrew a short distance and observed the animal whilst it recovered from immobilisation.

#### Biologging collars

Wild boar were fitted with Vertex Plus collars, produced by Vectronic Aerospace GmBH (https://www.vectronic-aerospace.com/, Berlin, Germany). These collars include a standard GPS module which was set to record GPS fixes at 30-minute intervals. The collars used were customised to carry a Daily Diary (DD, Wildbyte Technologies Inc., Swansea, Wales) unit, encased in a resin housing on the collar side (Figure S3). DD include a tri-axial accelerometer and a tri-axial magnetometer, and these were programmed to record at 10hz (that is, ten records per second). GPS positions were transmitted to data storage via GSM module every 3.5 hours, and DD data was stored on an on-board micro-SD card. This data was downloaded upon recovery of the collar after drop-off. Maximum collar weight did not exceed 1100 grams, which corresponds to acceptable collar:animal weight ratios for standard tracking devices deployed on animals (see Wilson *et al.* (2021)).

Table S1: Minimum and maximum observed weather and day length values for the study period.

| Variable | Minimum recorded | Maximum recorded |
|---|---|---|
| Temperature (°C) | -15.1 | 36.9 |
| Humidity (%) | 32.8 | 99.9 |
| Precipitation (mm) | 0 | 31.2 |
| Snow depth (cm) | 0 | 20 |
| Day length (hours) | 8.05 | 16.38 |

### Supplementary Results

Table S2: Maximal model (A) and reduced model (B) for total sleep time (TST); effects were removed starting with the least meaningful as judged by $P_x$. We report the median posterior; lower and upper 95% credible interval of posterior draws; percentage-across-zero ($P_x$); Gelman-Rubin convergence statistic (Rhat); and effective sample size (ESS). Model fixed effects shown in the upper panel and random effects and correlations shown in the lower panel. For year, 2019 is the reference level and for area Doupov is the reference level.

| (A) | Parameter | Median | Lower 95% CI | Upper 95% CI | $P_x$ | Rhat | ESS |
|---|---|---|---|---|---|---|---|
| Fixed Effects | Intercept | 14.16 | 11.06 | 17.38 | 0.00 | 1.00 | 2493.19 |
| | Sigma Intercept | 0.62 | 0.53 | 0.71 | 0.00 | 1.00 | 2307.71 |
| | Temperature | -0.55 | -0.75 | -0.34 | 0.00 | 1.00 | 2694.87 |
| | Humidity | -0.14 | -0.26 | -0.02 | 1.68 | 1.00 | 2810.23 |
| | Precipitation | 0.01 | -0.08 | 0.10 | 39.94 | 1.00 | 2807.78 |
| | Snow Depth | 0.09 | -0.05 | 0.23 | 10.39 | 1.00 | 2941.60 |
| | Day Length | -0.22 | -0.41 | -0.03 | 1.03 | 1.00 | 2387.04 |
| | Moonlight | -0.13 | -0.23 | -0.03 | 0.57 | 1.00 | 2736.90 |
| | Area (Kostolec) | 1.94 | 0.40 | 3.40 | 0.86 | 1.00 | 2155.42 |
| | Sex (Male) | 1.21 | -0.37 | 2.71 | 6.35 | 1.00 | 2855.19 |
| | Year (2020) | -3.46 | -4.88 | -2.04 | 0.00 | 1.00 | 2712.77 |
| | Year (2021) | -4.62 | -5.80 | -3.42 | 0.00 | 1.00 | 2603.46 |
| | Month (poly 1) | -39.05 | -82.92 | -0.53 | 2.36 | 1.00 | 2816.63 |
| | Month (poly 2) | -20.74 | -42.47 | -0.55 | 2.21 | 1.00 | 2699.30 |
| Random Effects | sd Intercept | 1.19 | 0.81 | 1.73 | 0.00 | 1.00 | 2798.19 |
| | sd Month (poly 1) | 94.14 | 69.21 | 131.79 | 0.00 | 1.00 | 2653.65 |
| | sd Month (poly 2) | 2.78 | 0.13 | 13.86 | 0.00 | 1.00 | 2581.22 |
| | sd Sigma Intercept | 0.24 | 0.17 | 0.33 | 0.00 | 1.00 | 2809.27 |
| | cor Intercept ~ Month (poly 1) | -0.13 | -0.52 | 0.34 | 28.44 | 1.00 | 2911.27 |
| | cor Intercept ~ Month (poly 2) | -0.02 | -0.82 | 0.79 | 48.18 | 1.00 | 2831.96 |
| | cor Month (poly 1) ~ Month (poly 2) | -0.03 | -0.81 | 0.80 | 47.97 | 1.00 | 2808.61 |
| | cor Intercept ~ Sigma Intercept | 0.06 | -0.38 | 0.49 | 40.36 | 1.00 | 2931.21 |
| | cor Month (poly 1) ~ Sigma Intercept | -0.21 | -0.55 | 0.20 | 14.74 | 1.00 | 2554.93 |
| | cor Month (poly 2) ~ Sigma Intercept | 0.21 | -0.73 | 0.86 | 33.69 | 1.00 | 1569.39 |

| (B) | Parameter | Median | Lower 95% CI | Upper 95% CI | $P_x$ | Rhat | ESS |
|---|---|---|---|---|---|---|---|
| **Fixed Effects** | Intercept | 14.39 | 11.42 | 17.50 | 0.00 | 1.00 | 2715.90 |
| | Sigma Intercept | 0.63 | 0.53 | 0.72 | 0.00 | 1.00 | 2753.83 |
| | Temperature | -0.55 | -0.76 | -0.36 | 0.00 | 1.00 | 2773.40 |
| | Humidity | -0.12 | -0.24 | -0.01 | 1.39 | 1.00 | 2676.19 |
| | Day length | -0.22 | -0.41 | -0.04 | 0.79 | 1.00 | 2788.60 |
| | Moonlight | -0.14 | -0.24 | -0.04 | 0.32 | 1.00 | 2702.39 |
| | Area (Kostelec) | 1.96 | 0.27 | 3.43 | 1.36 | 1.00 | 2769.93 |
| | Year (2020) | -3.34 | -4.88 | -1.90 | 0.00 | 1.00 | 2578.33 |
| | Year (2021) | -4.70 | -5.94 | -3.42 | 0.00 | 1.00 | 2713.47 |
| | Month (poly 1) | -39.31 | -82.81 | 2.38 | 3.28 | 1.00 | 2573.72 |
| | Month (poly 2) | -21.27 | -43.01 | -0.81 | 2.07 | 1.00 | 2751.39 |
| **Random Effects** | sd Intercept | 1.23 | 0.85 | 1.86 | 0.00 | 1.00 | 2801.09 |
| | sd Month (poly 1) | 96.26 | 70.16 | 134.07 | 0.00 | 1.00 | 3004.93 |
| | sd Month (poly 2) | 2.71 | 0.10 | 14.46 | 0.00 | 1.00 | 2734.37 |
| | sd Sigma Intercept | 0.24 | 0.17 | 0.33 | 0.00 | 1.00 | 2666.16 |
| | cor Intercept ~ Month (poly 1) | -0.15 | -0.53 | 0.30 | 25.48 | 1.00 | 2783.37 |
| | cor Intercept ~ Month (poly 2) | -0.02 | -0.80 | 0.79 | 48.54 | 1.00 | 2711.94 |
| | cor Month (poly 1) ~ Month (poly 2) | 0.02 | -0.80 | 0.80 | 48.75 | 1.00 | 2992.20 |
| | cor Intercept ~ Sigma Intercept | 0.11 | -0.32 | 0.52 | 31.19 | 1.00 | 2544.74 |
| | cor Month (poly 1) ~ Sigma Intercept | -0.20 | -0.55 | 0.19 | 16.45 | 1.00 | 3045.81 |
| | cor Month (poly 2) ~ Sigma Intercept | 0.18 | -0.75 | 0.86 | 36.72 | 1.00 | 1939.05 |

Table S3: Maximal model (A) and reduced model (B) for number of bouts per day. We report the median posterior; lower and upper 95% credible interval of posterior draws; percentage-across-zero ($P_x$); Gelman-Rubin convergence statistic (Rhat); and effective sample size (ESS). Model fixed effects shown in the upper panel and random effects and correlations shown in the lower panel. For year, 2019 is the reference level and for area Doupov is the reference level.

| (A) | Parameter | Median | Lower 95% CI | Upper 95% CI | $P_x$ | Rhat | ESS |
|---|---|---|---|---|---|---|---|
| Fixed Effects | Intercept | 21.54 | 10.00 | 33.25 | 0.00 | 1.00 | 2087.97 |
| | Sigma Intercept | 1.82 | 1.74 | 1.90 | 0.00 | 1.00 | 2533.20 |
| | Temperature | 1.35 | 0.60 | 2.08 | 0.04 | 1.00 | 2702.69 |
| | Humidity | -0.62 | -1.04 | -0.22 | 0.18 | 1.00 | 2970.47 |
| | Precipitation | -0.23 | -0.52 | 0.06 | 6.10 | 1.00 | 2745.68 |
| | Snow Depth | -0.40 | -0.83 | 0.00 | 2.61 | 1.00 | 2863.91 |
| | Day Length | 0.05 | -0.65 | 0.74 | 44.25 | 1.00 | 2272.40 |
| | Moonlight | -0.12 | -0.47 | 0.25 | 25.45 | 1.00 | 2581.73 |
| | Area (Kostolec) | -1.31 | -7.24 | 4.53 | 32.23 | 1.00 | 2734.01 |
| | Sex (Male) | -1.39 | -7.30 | 4.30 | 30.69 | 1.00 | 2821.85 |
| | Year (2020) | 2.63 | -2.20 | 7.92 | 14.28 | 1.00 | 2958.53 |
| | Year (2021) | 6.15 | 1.62 | 10.69 | 0.50 | 1.00 | 2802.60 |
| | Month (poly 1) | -96.38 | -252.73 | 59.01 | 10.60 | 1.00 | 2531.06 |
| | Month (poly 2) | 61.21 | -87.67 | 238.54 | 22.27 | 1.00 | 2649.21 |
| Random Effects | sd Intercept | 5.07 | 3.04 | 8.06 | 0.00 | 1.00 | 2750.18 |
| | sd Month (poly 1) | 331.34 | 216.54 | 491.06 | 0.00 | 1.00 | 2461.10 |
| | sd Month (poly 2) | 269.54 | 167.46 | 409.97 | 0.00 | 1.00 | 2420.36 |
| | sd Sigma Intercept | 0.19 | 0.14 | 0.26 | 0.00 | 1.00 | 2866.26 |
| | cor Intercept ~ Month (poly 1) | -0.18 | -0.67 | 0.35 | 25.45 | 1.00 | 2555.10 |
| | cor Intercept ~ Month (poly 2) | 0.00 | -0.59 | 0.48 | 49.96 | 1.00 | 2691.71 |
| | cor Month (poly 1) ~ Month (poly 2) | -0.64 | -0.87 | -0.18 | 0.50 | 1.00 | 2820.35 |
| | cor Intercept ~ Sigma Intercept | 0.50 | 0.03 | 0.82 | 1.86 | 1.00 | 2792.96 |
| | cor Month (poly 1) ~ Sigma Intercept | 0.00 | -0.39 | 0.40 | 49.46 | 1.00 | 2563.99 |
| | cor Month (poly 2) ~ Sigma Intercept | -0.44 | -0.74 | -0.03 | 1.82 | 1.00 | 2821.58 |

| (B) | Parameter | Median | Lower 95% CI | Upper 95% CI | $P_x$ | Rhat | ESS |
|---|---|---|---|---|---|---|---|
| **Fixed Effects** | Intercept | 21.04 | 17.58 | 24.52 | 0.00 | 1.00 | 2699.66 |
| | Sigma Intercept | 1.82 | 1.74 | 1.90 | 0.00 | 1.00 | 2756.25 |
| | Temperature | 1.31 | 0.59 | 2.05 | 0.00 | 1.00 | 2753.25 |
| | Humidity | -0.78 | -1.14 | -0.40 | 0.00 | 1.00 | 2735.22 |
| | Snow Depth | -0.44 | -0.84 | -0.03 | 1.82 | 1.00 | 2743.92 |
| | Year (2020) | 2.72 | -2.58 | 7.58 | 13.17 | 1.00 | 2741.28 |
| | Year (2021) | 6.14 | 1.72 | 10.44 | 0.68 | 1.00 | 2876.95 |
| | Month (poly 1) | -94.21 | -242.26 | 55.77 | 10.85 | 1.00 | 2916.89 |
| | Month (poly 2) | 66.31 | -64.64 | 228.00 | 18.27 | 1.00 | 2616.87 |
| **Random Effects** | sd Intercept | 4.78 | 2.97 | 7.35 | 0.00 | 1.00 | 2736.91 |
| | sd Month (poly 1) | 326.33 | 218.40 | 475.79 | 0.00 | 1.00 | 2762.93 |
| | sd Month (poly 2) | 268.41 | 168.43 | 404.78 | 0.00 | 1.00 | 2766.90 |
| | sd Sigma Intercept | 0.19 | 0.14 | 0.26 | 0.00 | 1.00 | 2870.26 |
| | cor Intercept ~ Month (poly 1) | -0.21 | -0.65 | 0.32 | 22.56 | 1.00 | 2798.74 |
| | cor Intercept ~ Month (poly 2) | -0.05 | -0.58 | 0.43 | 42.51 | 1.00 | 2703.75 |
| | cor Month (poly 1) ~ Month (poly 2) | -0.66 | -0.88 | -0.24 | 0.39 | 1.00 | 2738.60 |
| | cor Intercept ~ Sigma Intercept | 0.53 | 0.06 | 0.84 | 1.43 | 1.00 | 2584.01 |
| | cor Month (poly 1) ~ Sigma Intercept | 0.01 | -0.39 | 0.40 | 47.86 | 1.00 | 2804.12 |
| | cor Month (poly 2) ~ Sigma Intercept | -0.43 | -0.72 | -0.03 | 1.89 | 1.00 | 2796.39 |

Table S4: Maximal model (A) and reduced model (B) for the longest bout per day. We report the median posterior; lower and upper 95% credible interval of posterior draws; percentage-across-zero ($P_x$); Gelman-Rubin convergence statistic (Rhat); and effective sample size (ESS). Model fixed effects shown in the upper panel and random effects and correlations shown in the lower panel. For year, 2019 is the reference level and for area Doupov is the reference level.

| (A) | Parameter | Median | Lower 95% CI | Upper 95% CI | $P_x$ | Rhat | ESS |
|---|---|---|---|---|---|---|---|
| Fixed Effects | Intercept | 5.43 | 4.98 | 5.87 | 0.00 | 1.00 | 2045.03 |
| | Sigma Intercept | -0.86 | -0.93 | -0.78 | 0.00 | 1.00 | 1989.98 |
| | Temperature | -0.08 | -0.12 | -0.04 | 0.00 | 1.00 | 1982.71 |
| | Humidity | 0.00 | -0.03 | 0.03 | 49.98 | 1.00 | 2033.56 |
| | Precipitation | 0.02 | 0.00 | 0.04 | 1.65 | 1.00 | 2123.76 |
| | Snow Depth | 0.03 | 0.01 | 0.04 | 0.05 | 1.00 | 1956.83 |
| | Day Length | -0.02 | -0.05 | 0.01 | 6.80 | 1.00 | 2030.98 |
| | Moonlight | 0.00 | -0.02 | 0.02 | 45.53 | 1.00 | 1877.90 |
| | Area (Kostolec) | 0.09 | 0.00 | 0.19 | 2.70 | 1.00 | 2016.63 |
| | Sex (Male) | 0.09 | -0.01 | 0.19 | 4.15 | 1.00 | 1971.06 |
| | Year (2020) | -0.31 | -0.45 | -0.18 | 0.00 | 1.00 | 1893.41 |
| | Year (2021) | -0.76 | -0.85 | -0.67 | 0.00 | 1.00 | 1853.47 |
| | Month (poly 1) | -0.06 | -3.46 | 3.45 | 48.83 | 1.00 | 1805.96 |
| | Month (poly 2) | -0.53 | -5.14 | 3.87 | 40.68 | 1.00 | 2065.99 |
| Random Effects | sd Intercept | 0.02 | 0.00 | 0.09 | 0.00 | 1.00 | 1728.72 |
| | sd Month (poly 1) | 4.73 | 2.30 | 7.91 | 0.00 | 1.00 | 1935.31 |
| | sd Month (poly 2) | 4.70 | 2.13 | 7.78 | 0.00 | 1.00 | 1608.92 |
| | sd Sigma Intercept | 0.18 | 0.13 | 0.24 | 0.00 | 1.00 | 1981.84 |
| | cor Intercept ~ Month (poly 1) | 0.20 | -0.70 | 0.86 | 32.33 | 1.00 | 1798.07 |
| | cor Intercept ~ Month (poly 2) | -0.08 | -0.82 | 0.78 | 43.83 | 1.00 | 1798.74 |
| | cor Month (poly 1) ~ Month (poly 2) | -0.52 | -0.87 | 0.19 | 6.65 | 1.00 | 1953.07 |
| | cor Intercept ~ Sigma Intercept | -0.19 | -0.86 | 0.73 | 35.03 | 1.00 | 1176.75 |
| | cor Month (poly 1) ~ Sigma Intercept | 0.14 | -0.41 | 0.68 | 33.13 | 1.00 | 1670.43 |
| | cor Month (poly 2) ~ Sigma Intercept | 0.20 | -0.35 | 0.62 | 21.64 | 1.00 | 1780.56 |

| (B) | Parameter | Median | Lower 95% CI | Upper 95% CI | $P_x$ | Rhat | ESS |
|---|---|---|---|---|---|---|---|
| **Fixed Effects** | Intercept | 5.14 | 5.08 | 5.21 | 0.00 | 1.00 | 2795.84 |
| | Sigma Intercept | -0.86 | -0.94 | -0.79 | 0.00 | 1.00 | 2703.55 |
| | Temperature | -0.08 | -0.11 | -0.04 | 0.00 | 1.00 | 2742.27 |
| | Precipitation | 0.02 | 0.00 | 0.04 | 1.03 | 1.00 | 2639.46 |
| | Snow Depth | 0.03 | 0.01 | 0.04 | 0.04 | 1.00 | 2503.07 |
| | Year (2020) | -0.24 | -0.37 | -0.10 | 0.00 | 1.00 | 2586.80 |
| | Year (2021) | -0.71 | -0.80 | -0.61 | 0.00 | 1.00 | 2571.98 |
| | Month (poly 1) | 1.82 | -1.51 | 4.81 | 13.81 | 1.00 | 2477.36 |
| | Month (poly 2) | 1.94 | -1.82 | 5.21 | 15.74 | 1.00 | 2813.10 |
| **Random Effects** | sd Intercept | 0.04 | 0.00 | 0.11 | 0.00 | 1.00 | 2211.71 |
| | sd Month (poly 1) | 4.38 | 1.70 | 8.04 | 0.00 | 1.00 | 2474.84 |
| | sd Month (poly 2) | 4.94 | 1.88 | 8.25 | 0.00 | 1.00 | 2236.20 |
| | sd Sigma Intercept | 0.18 | 0.13 | 0.25 | 0.00 | 1.00 | 2774.01 |
| | cor Intercept ~ Month (poly 1) | 0.12 | -0.68 | 0.79 | 39.79 | 1.00 | 2514.29 |
| | cor Intercept ~ Month (poly 2) | -0.25 | -0.86 | 0.61 | 29.09 | 1.00 | 2158.64 |
| | cor Month (poly 1) ~ Month (poly 2) | -0.52 | -0.90 | 0.22 | 7.92 | 1.00 | 2683.77 |
| | cor Intercept ~ Sigma Intercept | -0.44 | -0.91 | 0.53 | 17.63 | 1.00 | 1715.19 |
| | cor Month (poly 1) ~ Sigma Intercept | 0.10 | -0.44 | 0.68 | 36.01 | 1.00 | 2520.02 |
| | cor Month (poly 2) ~ Sigma Intercept | 0.32 | -0.20 | 0.70 | 11.06 | 1.00 | 2614.09 |

### Supplementary Figures

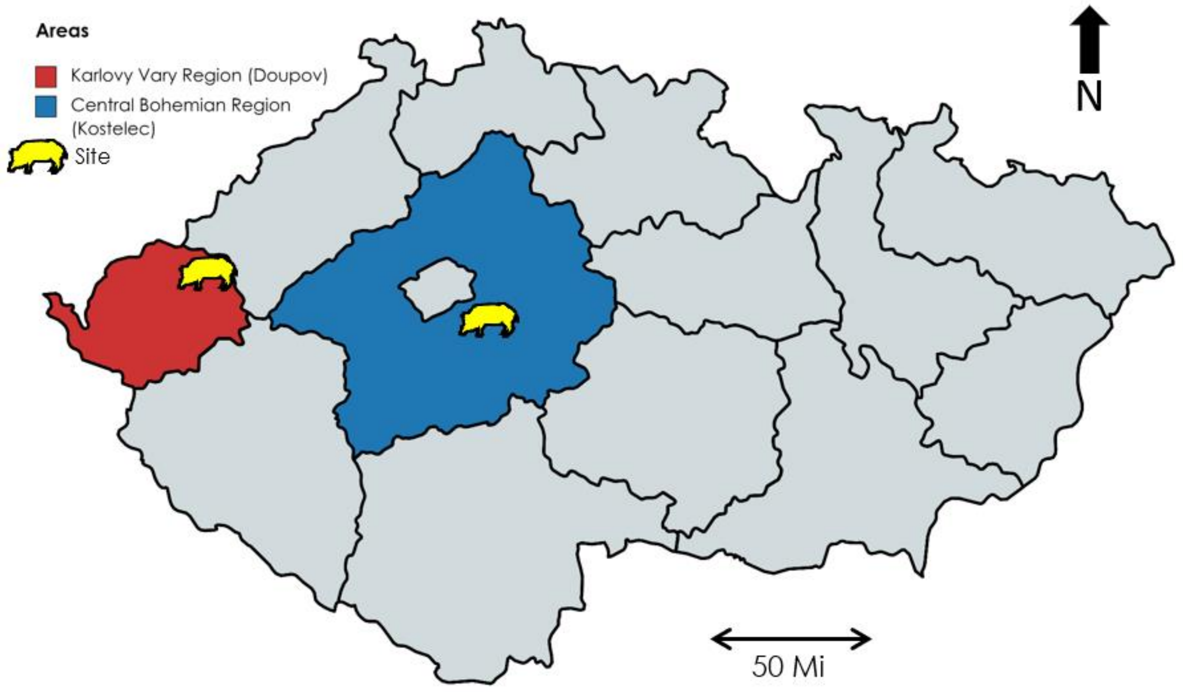


Figure S1: Map of Czech Republic showing locations of study sites (yellow) within their respective regions (blue and red). Scale for 50 miles shown lower right.

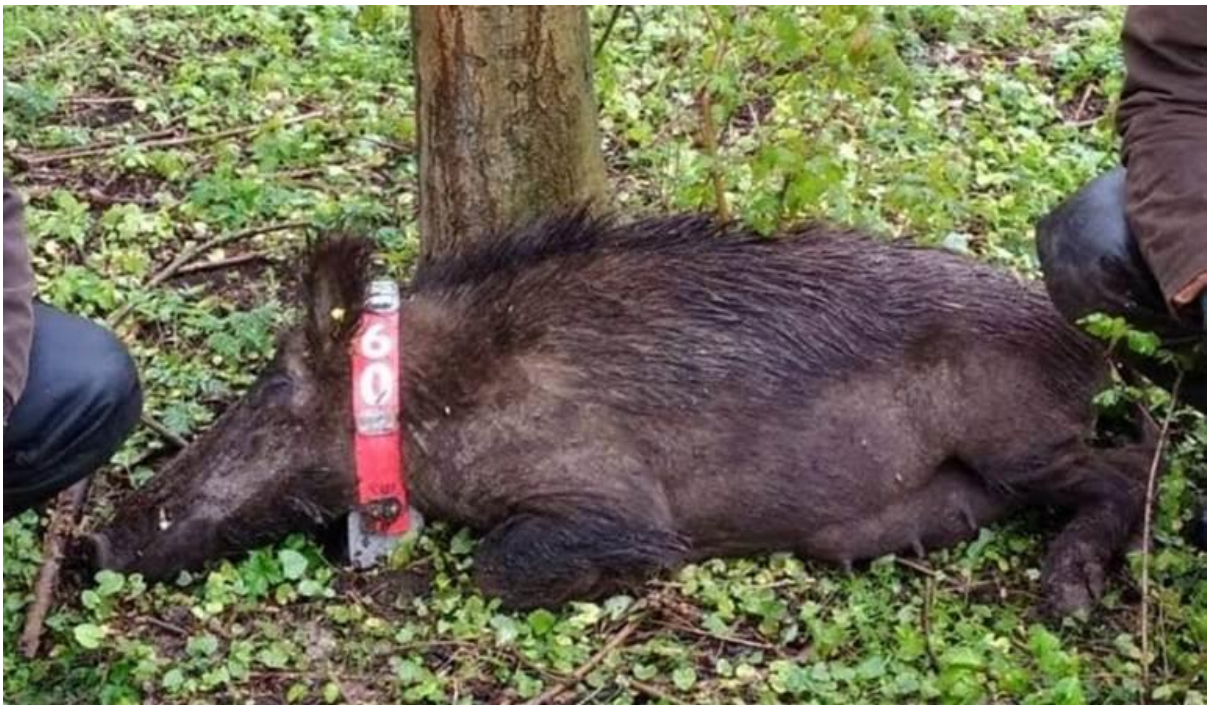


Figure S2: Wild boar after collar fitment, before recovery from immobilisation. DD is visible inside the resin casing under the number "60"

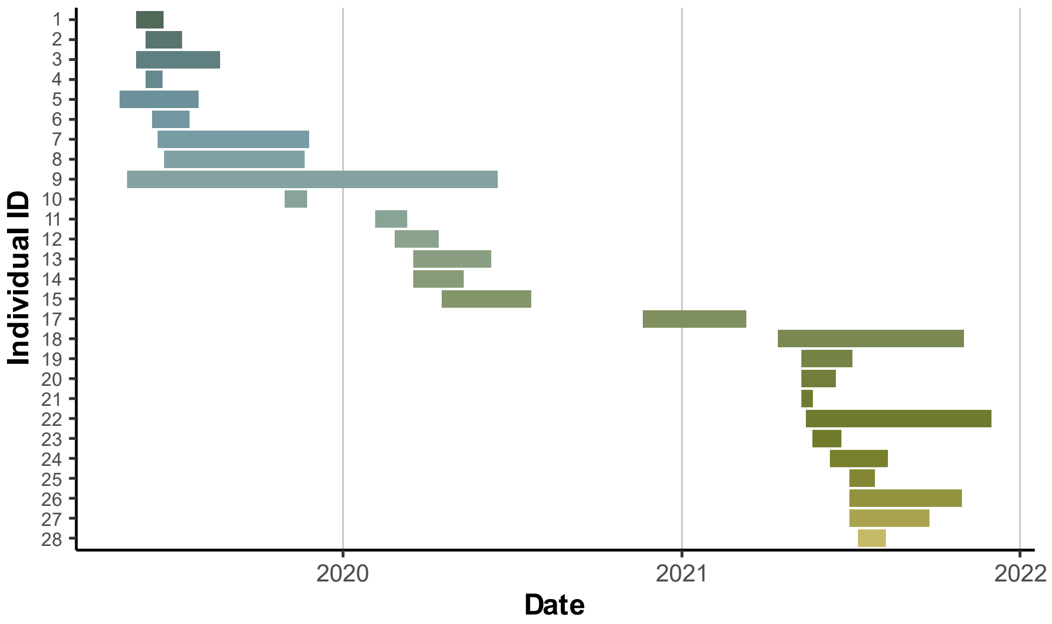


Figure S3: Recording duration by individual, starting on the 05/05/19 until the 01/12/21. Date shown on the x axis, where vertical gridlines indicate the start of a new year (31st December/1st of January).

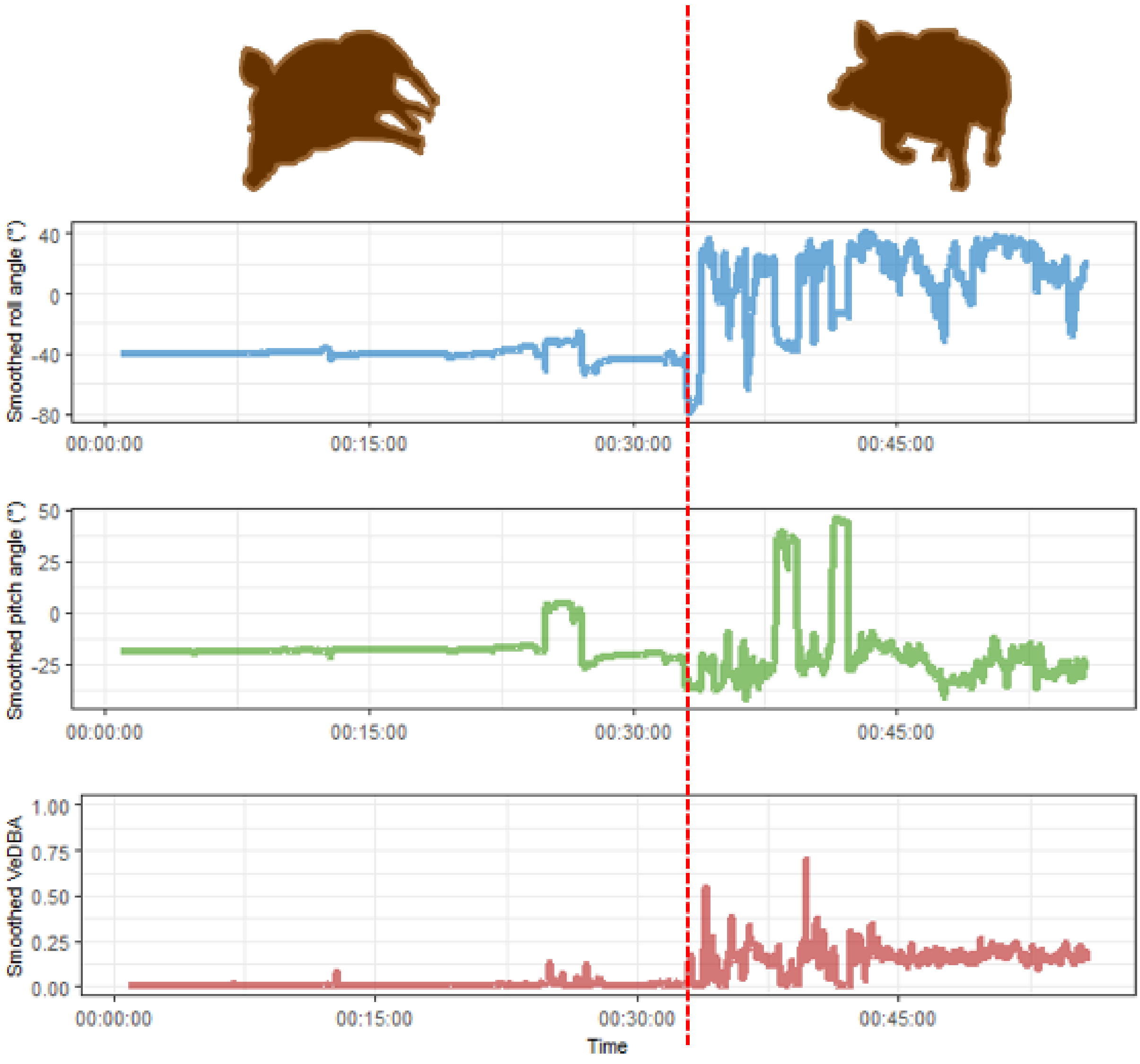


Figure S4: Lateral recumbency and the end of sleep bout. Visualisation of DD data showing changes in smoothed roll, pitch, and VeDBA values, corresponding to relevant behavioural types separated by red dashed lines, to identify the end of sleep bouts. Left, patterns typical of lateral recumbency with small in-sleep movement; right, patterns typical of highly active behaviour.

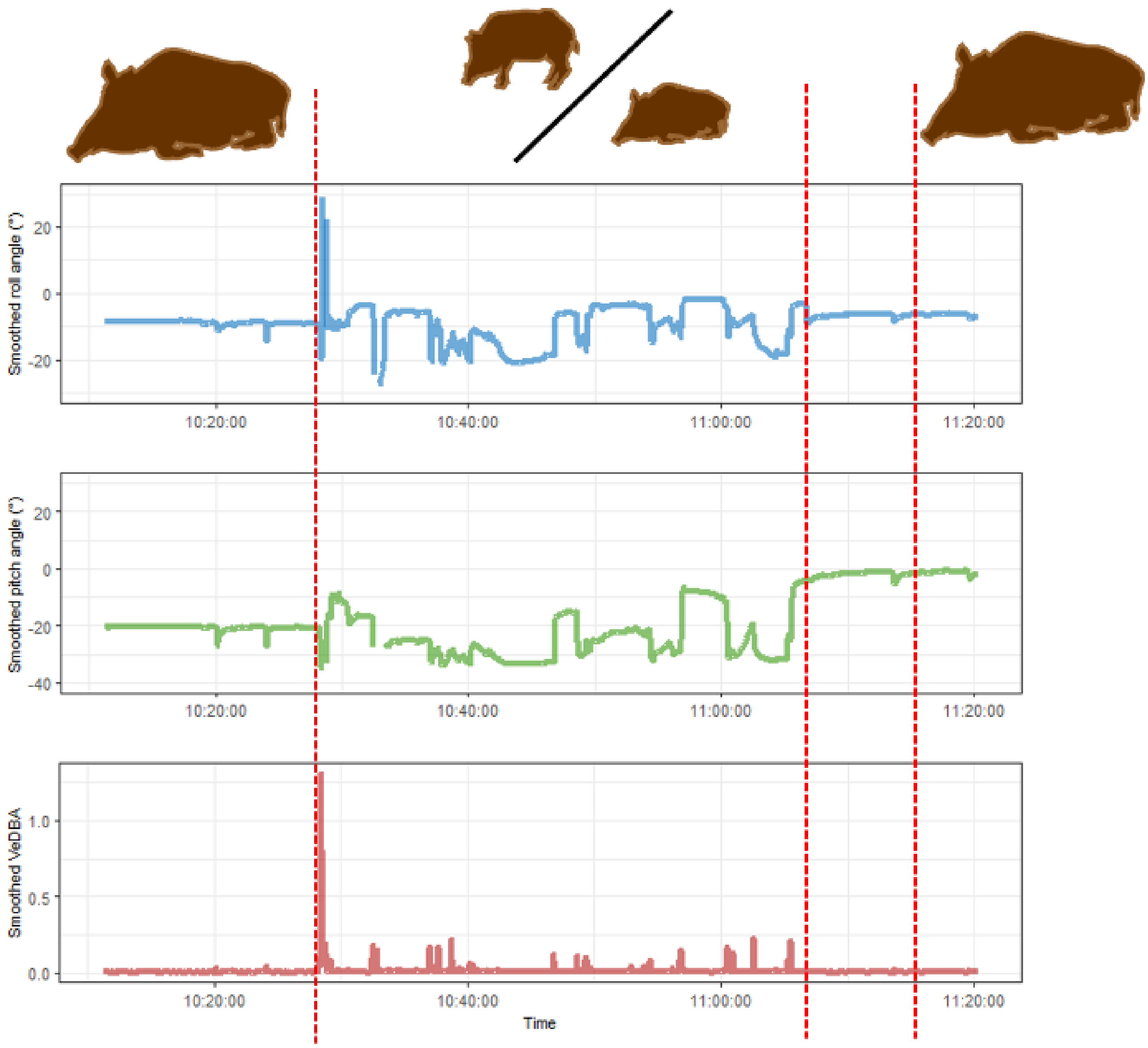


Figure S5: Sternal recumbency and wakeful rest. Visualisation of DD data showing changes in smoothed roll, pitch, and VeDBA values, corresponding to relevant behavioural types separated by red dashed lines. Left, sternal recumbency with small in-sleep movement (postural changes while sleeping); middle, wakeful rest and light activity; right, a return to sternal recumbency and sleep after transitional stage.

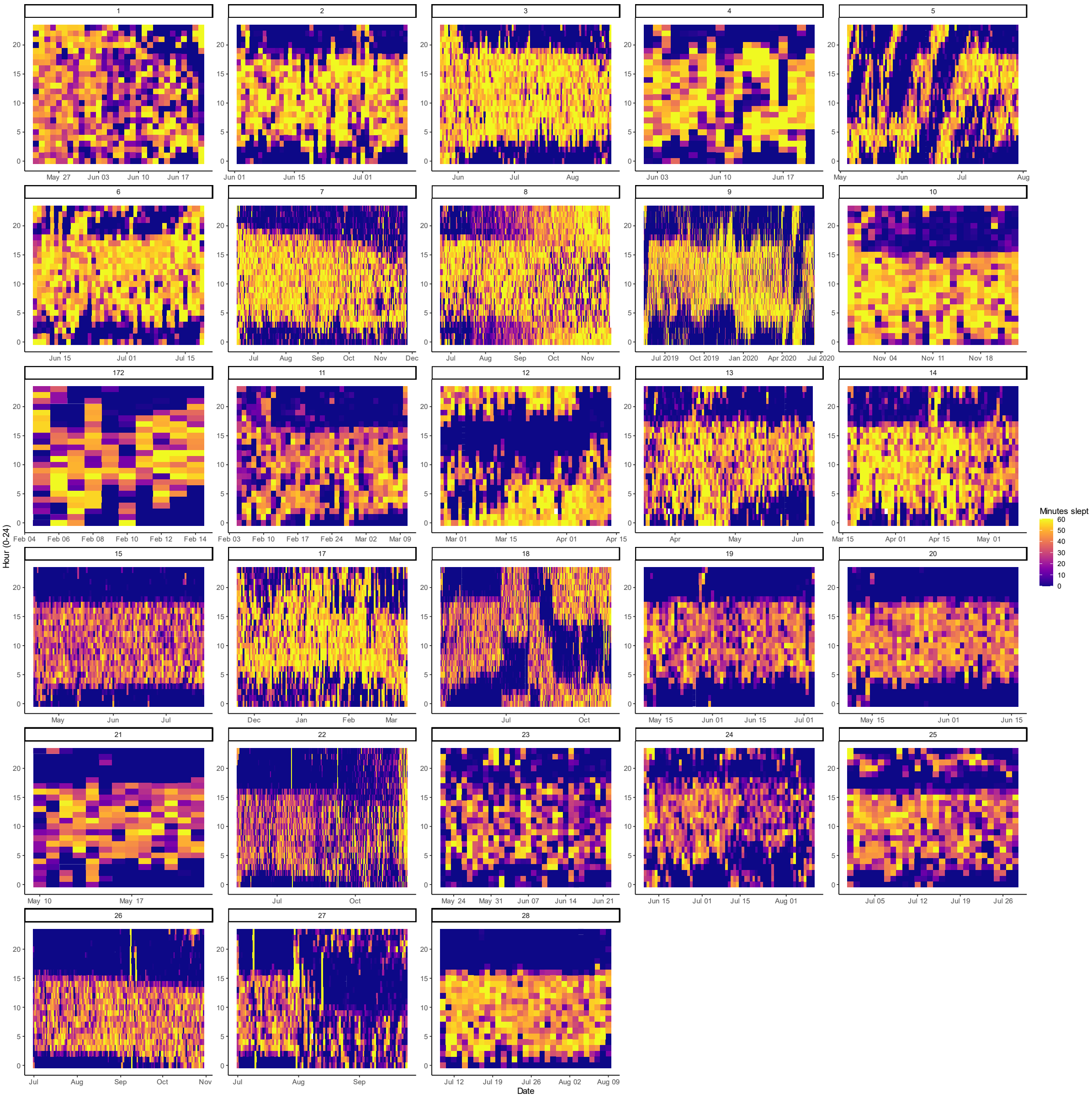


Figure S6: Minutes of sleep per hour, by day in each boar. Each column is one day split into 24 cells for each hour of the day, coloured by the number of minutes slept in that hour.
